# A miniaturized MR1 metabolite display system with native-like protein features

**DOI:** 10.64898/2026.04.13.718121

**Authors:** Photis Rotsides, Omkar Shinde, Julia N. Danon, Nikolaos G. Sgourakis

## Abstract

Major histocompatibility complex class I–related protein 1 (MR1) presents metabolite-derived antigens to mucosal-associated invariant T (MAIT) cells and other MR1-restricted T cells, playing a critical role in immune surveillance during infection and disease. Biochemical and structural studies of MR1 have been limited by the intrinsic instability of the molecule, which requires both ligand binding and association with beta-2-microglobulin (β_2_m) for proper folding and stability. Here, we adapt MR1 to the SMART protein platform to generate a minimalistic system for studying MR1 ligand presentation and T cell receptor (TCR) recognition. SMART-MR1 consists of the MR1 α_1_/α_2_ ligand-binding platform fused to a helical stabilizing domain that functionally replaces the α_3_ and β_2_m domains, resulting in a truncated protein that preserves the architecture of the antigen-binding groove. We show that SMART-MR1 can be efficiently produced recombinantly and retains the ability to bind chemically diverse classes of MR1 ligands. The reduced size of SMART-MR1 enables amide-based solution NMR experiments, and its simplified structure allows for ligand screening using fluorescence polarization. Importantly, SMART-MR1 maintains binding to the MAIT-derived A-F7 TCR, as confirmed by isothermal titration calorimetry. Finally, cryo-EM structural analysis of SMART-MR1/5-OP-RU bound to A-F7 revealed that ligand presentation and TCR recognition are nearly identical to those observed in native MR1. Together, these results establish SMART-MR1 as a minimal yet native-like system, expanding the experimental toolkit available for studying MR1 interactions and facilitating future efforts aimed at targeting MR1 pathways.

**Significance Statement:** MR1 is a highly conserved antigen-presenting molecule that enables T cells to detect metabolite signals from microbial infections and host metabolism. Despite its importance in immunity, mechanistic studies of MR1 have been limited by the instability of the native protein. We developed a simplified and stabilized version of MR1 that preserves the ligand-binding platform while eliminating structural elements that complicate biochemical analysis. This minimal system retains native-like antigen presentation and T cell receptor (TCR) recognition, while enabling experimental approaches that are difficult with full-length MR1. By lowering technical barriers to studying MR1–ligand and MR1–TCR interactions, this platform provides a versatile tool for exploring how antigens shape immune responses and for accelerating discovery of therapeutic strategies targeting MR1.

## Introduction

On the surface of all nucleated cells, class I major histocompatibility complex (MHC-I) molecules bind and present a repertoire of peptides derived from the endogenous proteome as a means of immune surveillance by CD8 T cells (1). In an analogous process, the non-classical MHC-I-related protein 1 (MR1) displays small molecule ligands derived from the endogenous and exogenous metabolome to Mucosal-associated invariant T (MAIT) cells and MR1-restricted T cells (2). The MR1 processing and presentation pathway begins with the assembly of a nascent MR1 heavy chain with the light chain beta-2 microglobulin (β_2_m) in the endoplasmic reticulum (ER) (3). The assembly process is facilitated by dedicated molecular chaperones: tapasin, which is restricted within the ER-resident peptide-loading complex (PLC), and the homologous, PLC-independent TAP-binding protein related (TAPBPR) (4, 5). However, unlike classical MHC-I, MR1 molecules are only transiently expressed at the cell surface and are rapidly internalized (3, 6). Under steady state conditions, nascent MR1 molecules are localized to the ER, vesicular pools, and endocytic compartments (7-9). These empty MR1 molecules are highly unstable and prone to aggregation. Therefore, MR1 folding and metabolite binding are cooperative processes. The availability of robust ligands triggers MR1 to egress out of the ER, and traffic to the cell surface (4, 10). Alternatively, partially folded MR1 can capture ligands on the cell surface or within vesicular pools, endocytic compartments, and phagosomes to orchestrate host responses (11). Finally, MR1 has a cytoplasmic tyrosine-based motif that is recognized by the endocytic adaptor protein 2 (AP2) complex. This interaction controls the kinetics of MR1 internalization from the cell surface, and minimizes recycling to define the duration of metabolite presentation (12). The importance of MR1 for human health is highlighted by the recent discovery that the human herpesviruses HSV-1 and CMV can disrupt MR1 expression through direct inhibition of the physiological ligand loading process (13).

Canonical MR1 ligands are derived from bacterial metabolite synthesis pathways (14, 15). The two prominent classes are lumazine/pyrimidine derivatives of riboflavin/vitamin B2 (16, 17) and pterins derived from folate/vitamin B9 (18, 19). While RL-6,7-DiMe, Ac-6-FP, and 5-OP-RU all upregulate MR1 surface expression, RL-6,7-DiMe and 5-OP-RU are agonists while 6-FP is antagonistic to T cells (14, 20). Other microbial MR1 ligands, such as 7,8-didemethyl-8-hydroxy-5-deazariboflavin (FO), photolumazines, hesperidin and riboflavin, exhibit pleiotropic properties (15). In contrast to the six polymorphic pockets in the peptide binding groove of MHC-I (A-F), the MR1 groove comprises only A’ and F’ pockets (18, 21). All MR1 ligands characterized to date bind exclusively to the A’ pocket rather than the much shallower MR1 F’ pocket (20). It has been proposed that the formation of a Schiff base between the conserved K43 buried in the A’ pocket and metabolites promotes stable MR1 folding (3, 18). However, several potent metabolites have been identified which associate with MR1 in a non-covalent manner but still upregulate surface expression levels (14, 22, 23). Consistent with this observation, MR1 K43A mutants can still bind ligands, albeit with decreased thermal stabilities (19, 24), suggesting that K43 Schiff base formation may not be the only determinant for MR1 egress out of the ER, and that both covalent and non-covalent ligands can form stable complexes with MR1. These knowledge gaps preclude a detailed understanding of what defines a physiologically relevant repertoire of ligands from the cellular metabolome (15).

Previous studies aiming to stabilize the MR1 structure for recombinant protein production and ligand screening applications have used either a chimeric approach where human α_1_/α_2_ platform is coupled to bovine α_3_/β_2_m domains (hpMR1)(25), or, more recently an “open” MR1 format, with an engineered disulfide bond at the heavy and light chain interface (26). We have recently developed SMART MHC proteins, which consist of a single-chain α_1_/α_2_ binding groove with the α_3_ and β_2_m domains replaced by a helical stabilizing domain (27). Here, we leverage the SMART platform to create a stable minimalistic system for studying MR1 *in vitro*. We demonstrate that SMART-MR1 can capture different classes of known ligands and maintains binding to cognate T cell receptors (TCRs), while also offering new avenues for studying MR1 interactions by solution NMR. We also solve the three-dimensional structure of SMART-MR1 in complex with A-F7, illustrating that SMART-MR1 presents ligands and binds to TCRs similarly to native MR1. Our results demonstrate that SMART-MR1 is a miniaturized system with native-like properties, expanding the potential for studying MR1 using standard biochemical and structural techniques.

## Results

### Design and purification of SMART-MR1

To determine whether MR1 is a suitable candidate for adapting it to the SMART system, we compared the interface formed by β_2_m and the α_1_/α_2_ “platform” between the structures of human metabolite-bound MR1 (PDB 6PUC) and peptide-bound HLA-A*02:01 (A02, PDB 9SKO). Despite sharing limited (43.6%) overall sequence homology, the α_1_/α_2_ domains of MR1 and A02 bear considerable structural similarity (Fig. 1A). In both structures, the interface is formed by polar and hydrophobic contacts between key β_2_m and α_1_/α_2_ residues that are conserved between A02 and MR1. Specifically, W60 from β_2_m is nestled within a hydrophobic pocket formed by the conserved residue pair M98/95 and F8 under the α_1_/α_2_ β-sheet, and this interaction plays a central role in both structures. The W60 ring forms a hydrogen bond with D122/118 in A02/MR1, further stabilizing the β_2_m/heavy chain interface. Indeed, a key tryptophan at the analogous position in the interface was shown to be favorable when screening for different SMART MHC stabilizing domains (27). Additional stabilizing polar contacts are formed between β_2_m residues R3 and D53 with the analogous residues R48/46 and G120/116 in A02/MR1. This striking degree of structural similarity of the β_2_m/heavy chain interface in MR1 and A02 suggests that the SMART MHC stabilizing domain can also form the necessary contacts required to stabilize the α_1_/α_2_ MR1 platform. To this end, we generated a SMART-MR1 construct by joining the N-terminus of the MR1 α_1_/α_2_ domain to the C-terminus of the SMART MHC stabilizing domain via a rigid linker. Using AlphaFold3, we generated a structural model of SMART-MR1 and found that it closely recapitulated the features of the SMART MHC X-ray structure. In this model, the stabilizing domain formed a network of favorable contacts with the MR1 α_1_/α_2_ platform. Notably W17 of the SMART-MR1 occupied the hydrophobic pocket under the α_1_/α_2_ β-sheet, playing an equivalent role to W60 from β_2_m in the native MR1 structure (Fig. 1B).

**Figure 1.**
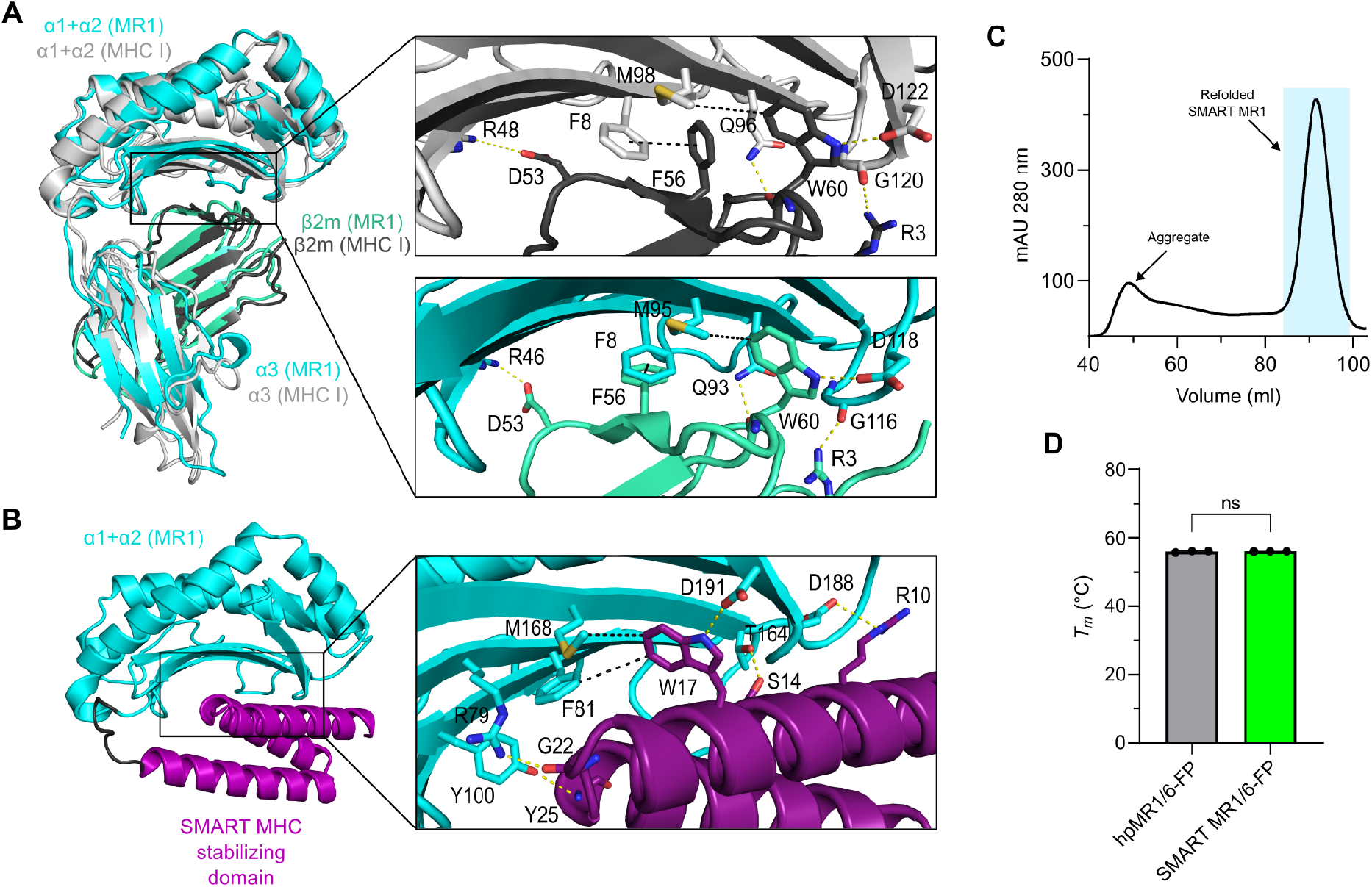
Design and purification of SMART-MR1. **A**. Crystal structure of human MR1/β_2_m (PDB 6PUC) superimposed onto the crystal structure of human HLA-A*02:01/β_2_m (PDB 9SKO). Insets illustrate key interactions that stabilize the β_2_m interface with the α_1_/α_2_ domains of HLA-A*02:01/MR1. Polar and hydrophobic contacts are indicated by yellow and black dashes, respectively. **B**. AlphaFold model of SMART-MR1 using the α_1_/α_2_ domain of MR1 and the stabilizing domain of SMART MHC. The inset displays contacts between the stabilizing domain and the α_1_/α_2_ domain of MR1. The rigid linker connecting the two domains is colored black. **C**. Size exclusion chromatography (SEC) trace of SMART-MR1 refolded in the presence of 6-FP. **D**. Melting temperatures (*T*_m_, °C) obtained from DSF of SMART-MR1 and hpMR1/bβ_2_m refolded in the presence of 6-FP. Data are mean ± s.d. for *n* = 3 technical replicates. Unpaired t-test was performed to obtain a P value of 0.6713, P > 0.05 (ns).

We next sought to express and purify SMART-MR1 from *E. coli* lysates. Our previous work has shown that, while SMART MHC can be expressed in soluble, peptide-receptive form, refolding of the same protein in the presence of excess peptide results in enhanced native structural features and improved affinity for cognate T cell receptors (TCRs) (27). These results suggest that refolded SMART proteins confer superior functionality. We therefore chose to refold SMART-MR1 from *E. coli* inclusion bodies using standard procedures (25, 28). Purification of SMART-MR1 refolded in the presence of the covalent ligand 6-FP by size-exclusion chromatography (SEC) resulted in a monodisperse peak at the expected molecular weight (Fig.1C and *SI Appendix*, Fig. S1A). To evaluate the quality of the purified SMART-MR1/6-FP complex we compared its thermal stability to full-length hpMR1/6-FP using differential scanning fluorimetry (DSF). The thermal stability of MR1 complexes is highly dependent on the occupancy a ligand within the A’ pocket (5), thus MR1 unfolding is coupled with dissociation of the ligand from the protein. The melting temperature (*T*_*m*_, °C) of SMART-MR1/6-FP was nearly identical to hpMR1/6-FP (56.0 vs 55.9 °C), indicating that SMART-MR1 can bind ligands and present them in a native-like manner (Fig. 1D and *SI Appendix*, Fig. S1B). Together, these results demonstrate that the SMART stabilizing domain can bolster the ligand-binding α_1_/α_2_ platform of MR1 effectively serving as a functional surrogate of the α_3_ and β_2_m domains.

### Engineered SMART-MR1 can accommodate several classes of known ligands

MR1 is capable of binding to multiple chemically distinct classes of small molecule metabolites. To explore whether SMART-MR1 can capture these diverse ligands, we refolded SMART-MR1 in the absence (empty) or presence of vitamin B9 metabolites (Ac-6-FP, 6-FP), vitamin B2 metabolites (5-OP-RU), and drug or drug-like molecules (3FSA, 5FSA) (Fig. 2A). For all ligands tested, refolding yields improved compared to refolding in the absence of ligand (Fig. 2B and *SI Appendix*, Fig. S2 A and C). Moreover, *T*_*m*_ values could only be determined when SMART-MR1 was refolded in the presence of a ligand (Fig. 2C and *SI Appendix*, Fig. S2B and C), likely due to the binding of the hydrophobic dye to the groove of empty SMART-MR1. *T*_*m*_ values also correlated with refolding yields (R^2^ = 0.503, Fig. 2D). These results indicate that SMART-MR1 binds to different classes of known MR1 ligands, and that, like native MR1, its protein stability is highly dependent upon the presence of bound ligand.

**Figure 2.**
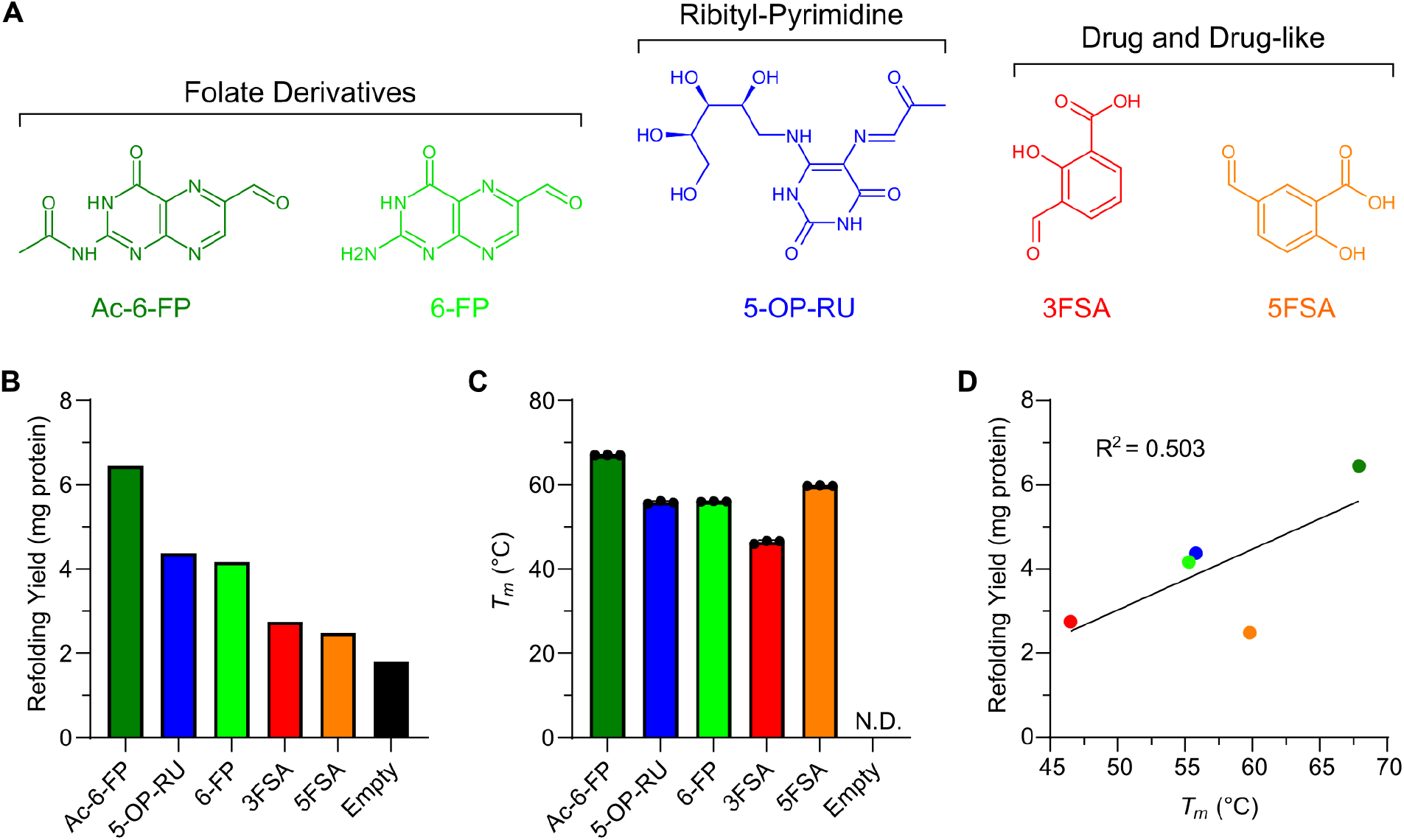
Ligand dependence of *in vitro* refolding and thermal stability of SMART-MR1 complexes. **A**. Chemical structures of ligands tested. **B**. Refolding yields (total mg protein) from SEC purification of SMART-MR1 refolded in the presence of ligands. **C**. *T*_m_ values obtained from DSF experiments of SMART-MR1 refolded in the presence of ligands. Data are mean ± s.d. for *n* = 3 technical replicates. **D**. Correlation plot of *T*_m_ versus refolding yield with the coefficient of determination from linear regression (R^2^) shown.

### SMART-MR1 is properly conformed and binds the A-F7 TCR with high affinity

To further characterize the structural and biochemical properties of SMART-MR1, and to explore potential applications of our system, we applied solution-based biophysical techniques. A major advantage of SMART-MR1 is that it drastically reduces the size of the complex compared to native MR1 (43 vs 29 kDa). This reduction in size mitigates relaxation losses during NMR experiments, allowing for facile backbone assignments and characterization of interactions with small molecules, immunoreceptors and molecular chaperones, whereas the study of WT MR1 necessitates more elaborate sample preparation with site-selective isotopic labelling (5). To test the feasibility of amide-based NMR experiments, we refolded isotopically labeled SMART-MR1 with ^2^H, ^13^C, and ^15^N in complex with 6-FP, and performed 2D TROSY experiments (Fig. 3A). The 2D TROSY spectrum was well dispersed, as expected for a properly conformed globular protein of 29 kDa, indicating that residue-level information on dynamics and interactions with different binding partners can be obtained using standard solution NMR methods.

**Figure 3.**
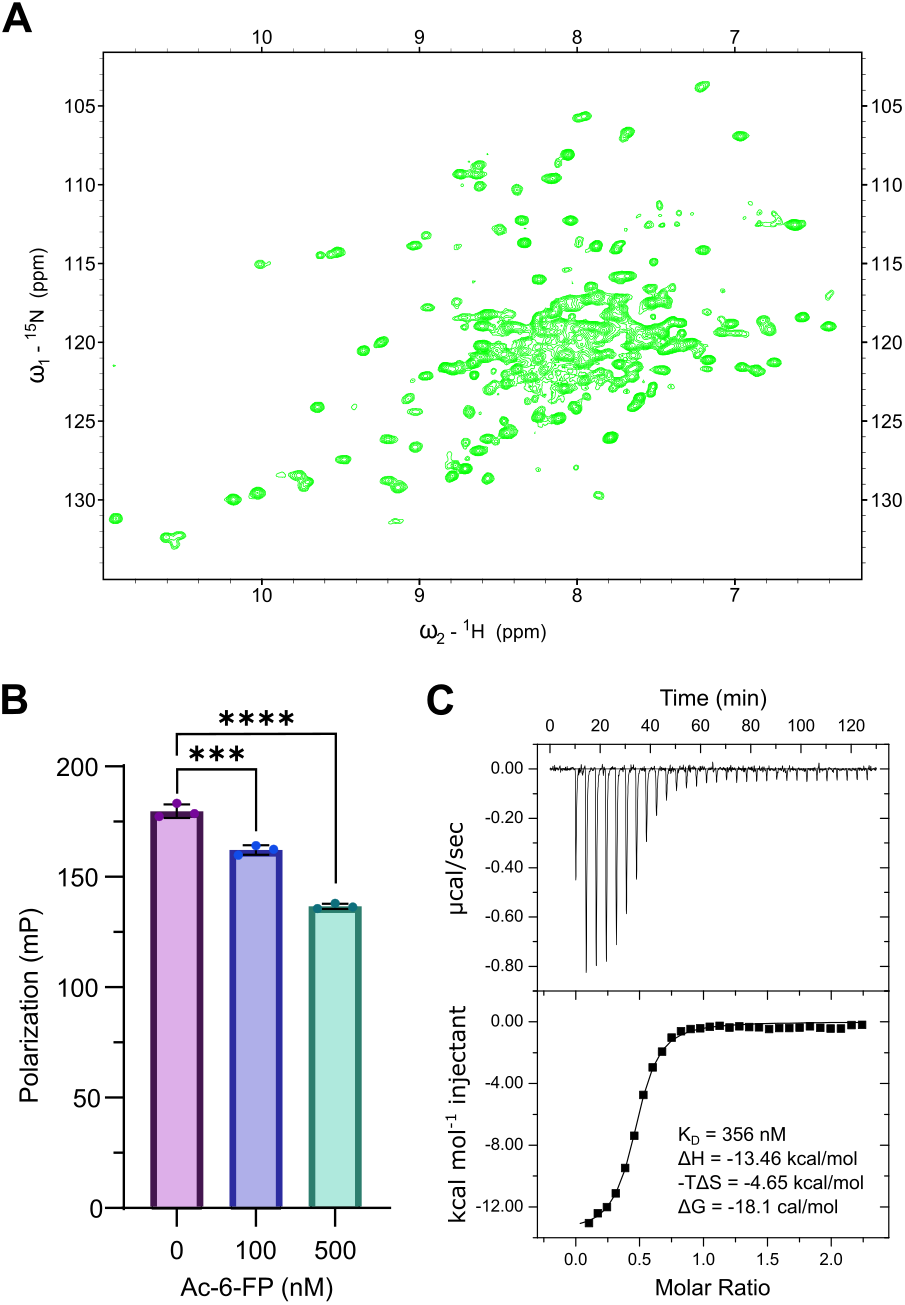
Solution biophysics of SMART-MR1. **A**. 2D ^1^H-^15^N TROSY of 650 μM SMART-MR1/6-FP recorded at a ^1^H field of 800 MHz at 25 °C. **B**. Competitive binding of TAMRA-labelled JYM20 to SMART-MR1/DCF as a function of increasing Ac-6-FP concentration, measured by fluorescence polarization. Data are mean ± s.d. for *n* = 3 technical replicates. One-way ANOVA was performed relative to conditions without Ac-6-FP, P < 0.0002(***), and P < 0.0001(****). **C**. ITC data titrating 200 μM A-F7 TCR into a sample containing 20 μM SMART-MR1/5-OP-RU. Black line is the fit of the isotherm with the K_D_ and thermodynamic values determined using a 1-site binding model (n = 1).

We next sought to determine whether SMART-MR1 can be used as a tool to screen for ligand binding. We refolded SMART-MR1 with the non-covalent ligand DCF and used a fluorescent 5-OP-RU analog (JYM20) (4) to perform competition experiments in the presence of a second ligand using fluorescence polarization (FP)(Fig. 3B). Upon incubation of SMART-MR1/DCF with JYM20 for 72 hours at room temperature, we detected an increase in the FP of JYM20 as it transitions into a high MW protein complex. Addition of a competitive ligand (Ac-6-FP) to the sample caused a significant reduction of FP in a dose dependent manner.

Next, we tested binding of SMART-MR1 to an established cognate TCR using isothermal titration calorimetry (ITC) (Fig. 3C). A-F7 is a MAIT cell-derived TCR that specifically binds to MR1 molecules presenting 5-OP-RU. A-F7 binding to SMART-MR1/5-OP-RU was exothermic, driven by a high enthalpy change of -13.5 kcal/mol, and showed a dissociation constant (K_D_) of 356 nM, as measured by ITC. This result could suggest an affinity enhancement for SMART-MR1 relative to values reported previously for A-F7 binding to native MR1 (K_D_ of 1.1 uM), albeit these experiments were performed using Surface Plasmon Resonance (29). These results demonstrate that SMART-MR1 provides a system for high-resolution solution biophysics by NMR, high-throughput ligand screening, and TCR binding studies.

### Cryo-EM structure of the SMART-MR1/A-F7 TCR complex reveals a native antigen recognition mechanism

To determine whether SMART-MR1 presents ligands and binds to TCRs in a native-like manner, SEC-purified SMART-MR1/5-OP-RU/A-F7 TCR complex was used to prepare grids for cryogenic electron microscopy (cryo-EM). To overcome the potential preferred orientation issue of the complex during vitrification, 0.01% fluorinated octyl-maltoside was added to the sample prior to grid preparation. Micrographs from cryo-EM data collection revealed an even distribution of particles and 2D classes showed different orientations of the complex. The data processing resulted in a nearly complete cryo-EM map (3.08 Å) of the complex with well-defined densities that allowed unambiguous register of the protein sequence and 5-OP-RU ligand (*SI Appendix*, Fig. S3 and Table S1).

The cryo-EM structure of SMART-MR1 displayed remarkable agreement with the predicted AlphaFold3 model and the native MR1 crystal structure (Fig. 4A). W17 of SMART-MR1 adopts a similar conformation to W60 of β_2_m, supporting our prediction that a Trp at this position is key for stabilizing the underside of the MR1 α_1_/α_2_ domain. We observed a continuous density connecting K116 of SMART-MR1 (analogous to K43 in native MR1) to 5-OP-RU, confirming that SMART-MR1 covalently captures ligands through the formation of a Schiff base (*SI Appendix*, Fig. S4A). The conformation of 5-OP-RU and its interactions within the binding groove remain conserved across the SMART-MR1 and native MR1 structures (Fig. 4B and C). The uracil ring of 5-OP-RU is burrowed within a hydrophobic pocket formed by Y80/W142 and is further stabilized by hydrogen bonds to R82 and S97. The hydroxyl groups of the ribityl chain form a network of polar interactions with R82, R167, Y225, Q226, and W229.

**Figure 4.**
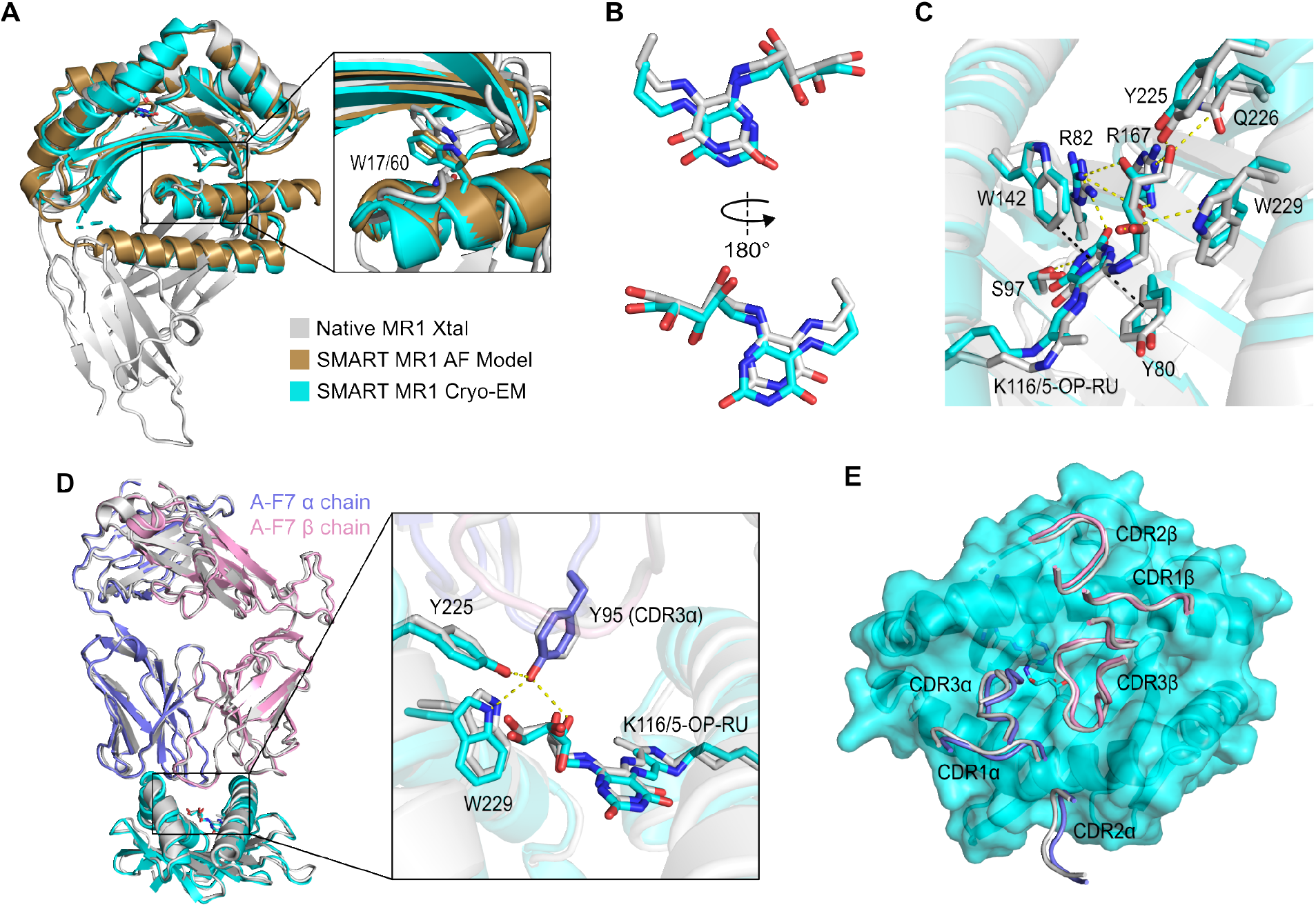
Cryo-EM structure of SMART-MR1/5-OP-RU bound to A-F7 TCR. **A**. Cryo-EM structure of SMART-MR1/5-OP-RU superimposed onto the AlphaFold model of SMART-MR1 and the crystal structure of human MR1/hβ_2_m-5-OP-RU (PDB 6PUC). The inset shows the sidechain of W17 of SMART-MR1 and W60 of hβ_2_m. **B**. Conformation of 5-OP-RU in the SMART-MR1 Cryo-EM structure superimposed onto 5-OP-RU from the native MR1 crystal structure. **C**. Close up of 5-OP-RU within the ligand binding groove of native and SMART-MR1. Polar and hydrophobic contacts are indicated by yellow and black dashes, respectively. Residues are colored as in panel A. **D**. Cryo-EM structure of A-F7 TCR bound to the α_1_/α_2_ domain of SMART-MR1/5-OP-RU (cyan) superimposed onto the native MR1/5-OP-RU-A-F7 crystal structure (grey). The inset highlights key interactions between the TCR and the ligand as well as residues within the MR1 groove. **E**. Top view of CDR loop interactions with the surface of SMART-MR1/5-OP-RU. CDR loops are shown as tubes and are colored as in panel D.

We also observed that the mechanism of 5-OP-RU recognition by A-F7 is nearly identical across the SMART and native MR1 structures. Y95 of the A-F7 complementarity determining region (CDR) 3α reaches into the MR1 binding groove and forms polar contacts with Y225 and W229 in addition to the ribityl group of 5-OP-RU (Fig. 4D). Furthermore, all six CDR loops of A-F7 dock onto the SMART-MR1 binding groove similarly to native MR1 (Fig. 4E). CDR loops 1α and 2α form contacts with the α_1_helix of SMART-MR1, whereas CDR loops 1β and 2β interact with the α_2_ helix. CDR loops 3α and 3β are positioned centrally above the MR1 binding groove, while only CDR3α is in a favorable position to extend into the A’ pocket to make direct contact with the ligand. Together, these structural results demonstrate that SMART-MR1 presents ligands and binds to TCRs in a manner nearly indistinguishable from native MR1.

## Discussion

Biochemical characterization of native MR1 presents challenges due to the fact that its stability is coupled to the presence of a ligand within the binding groove, in addition to association with β_2_m. Here, we demonstrate that the α_1_/α_2_ metabolite-presenting platform of MR1 can be stabilized through interactions with a synthetic protein domain, mitigating any concerns caused by β_2_m dissociation while maintaining binding to known ligands. Our biophysical data confirm that SMART-MR1 is stable and properly conformed, and that SMART-MR1 binds to cognate TCRs with high affinity. Our structural results demonstrate SMART-MR1 presents ligands in a native-like manner, and that TCRs bind to SMART and native MR1 through identical interactions.

Identifying new ligands and TCRs is critical for understanding MR1 biology in the context of cancer and infectious disease and for developing immunotherapies (22, 29-35). SMART-MR1 provides a platform for rapid exploration of ligand and TCR interactions by various orthogonal techniques. Screening ligands is possible by using SMART-MR1 for high-throughput competitive FP assays. The reduced size and high expression yield of SMART-MR1 allows for mapping of interactions with TCRs by NMR, as illustrated by our recent work using SMART MHC (36). As shown by our current structural determination of the complex with A-F7, SMART MR1 provides an attractive system for high-resolution structural studies of metabolite recognition by MAIT cell receptors using cryo-electron microscopy. Finally, while our work focuses on the MR1*01 allele which has a frequency 71% in humans (37), expanding SMART-MR1 to include less frequent alleles can help to determine the effect of polymorphisms on ligand binding and TCR recognition. In summary, SMART-MR1 offers a streamlined and adaptable system that maintains the functional features of MR1, while enabling a wide range of biochemical, structural, and screening approaches for characterizing emerging MR1-restricted antigens relevant to MAIT cell biology, and human disease.

## Materials and Methods

### MR1 ligands

MR1 ligands were purchased from the following suppliers: Ac-6-FP (Cayman Chemical no. 23303), 6-FP (Cayman Chemical no. 14247), 3FSA (3-formylsalicylic acid) (Acros Organics no. 377070250), 5FSA (5-formylsalicylic acid) (Millipore Sigma no. F17601), DCF (Millipore Sigma no. D6899). 5-OP-RU was synthesized in-house from 5-N-RU, which was generously provided by E. Adams (University of Chicago). Chemical structures were prepared using ChemSketch (ACD/Labs).

### Protein expression, refolding, and purification

Codon optimized DNA encoding SMART-MR1, hpMR1, or bovine β2m was transformed into BL21(DE3) *E. coli* (New England Biolabs). Proteins were expressed in autoinducing media at 37 °C overnight (38). Inclusion bodies were isolated and solubilized using guanidine hydrochloride (39). For *in vitro* refolding of SMART-MR1, 60 mg of solubilized inclusion body was slowly diluted into 300 mL of refolding buffer (0.4 M arginine–HCl, 2 mM EDTA, 4.9 mM reduced L-glutathione, 0.57 mM oxidized L-glutathione, 100 mM Tris pH 8.0) at 4 °C while stirring in the absence (empty) or presence of 2 mg of ligand. For hpMR1 refolding, a 50 mg mixture of hpMR1:bβ_2_m at a 1:1 molar ratio was slowly diluted into 1 L of refolding buffer containing 2 mg ligand. Refoldings proceeded at 4 °C for four days without stirring. Solutions were then dialyzed into buffer containing 150 mM NaCl, 25 mM Tris pH 8.0 overnight. The refolding mixtures were concentrated using Pellicon Single-Pass Tangential Flow Filtration (Millipore Sigma) and Amicon ultra centrifugal filters (Millipore Sigma). Purification of refolded MR1 proteins was performed by size-exclusion chromatography using a HiLoad 16/600 Superdex 200 pg column at 1 mL/min in running buffer (150 mM NaCl, 25 mM Tris pH 8.0).

### Differential scanning fluorimetry

DSF was used to assess the thermal stabilities of refolded MR1 molecules. 10 μM protein was mixed with 10× SYPRO Orange dye (Invitrogen) in a buffer of 150 mM NaCl and 20 mM sodium phosphate pH 7.4. 20 μL samples were loaded into MicroAmp Optical 384-well plate and the experiment was performed on a QuantStudio 5 real-time PCR machine with excitation and emission wavelengths set to 470 nm and 569 nm. The temperature was incrementally increased at a rate of 1 °C per minute between 25 and 95 °C. Data analysis and fitting were performed in GraphPad Prism v9.

### Nuclear magnetic resonance

To generate isotopically-labeled SMART-MR1, proteins were expressed in M9 media containing ^2^H_2_O, 3 g/L [^2^H, ^13^C] glucose, and 1 g/L of ^15^NH_4_Cl. Protein expression was induced with 1 mM IPTG at OD_600_=0.6 followed by shaking at 37 °C for 5 hours. Inclusion bodies and refolding mixtures were prepared as described above. SEC purification of triple-labeled protein was performed in NMR buffer (100 mM NaCl, 20 mM sodium phosphate, pH 7.2). NMR samples were prepared with 650 μM protein and 5% ^2^H_2_O. ^1^H-^15^N TROSY HSQC spectra were collected at 800 MHz ^1^H magnetic field. The data were processed in NMRPipe (40) and analyzed in POKY (41).

### Fluorescence polarization

The association of a TAMRA-labeled, fluorescent 5-OP-RU analog (JYM20) (4) to SMART-MR1/DCF was monitored using fluorescence polarization (FP). 10 nM JYM20 was incubated with 10 μM SMART-MR1/DCF in the absence or presence of increasing amounts of Ac-6-FP for 72 hours at room temperature in the dark. To determine the optimal incubation conditions for ligand binding, several association assays were performed at room temperature and in 4° C across various incubation times. We found that 48 hours was not sufficiently long enough to facilitate complete association and that at 96 hours a degree of protein degradation had occurred. Additionally, experiments performed at 4° C did not reach full equilibrium, even after 72 hours. Reactions performed at room temperature for 72 hours were able to both reach equilibrium and avoid degradation, a finding consistent across the literature (42). Excitation and emission values used to detect the fluorescence of JYM20 were 531 and 595 nm on a SpectraMax iD5 plate reader. Raw parallel (I_II_) and perpendicular emission intensities (I_⊥_) were collected and converted to polarization (mP) values using the equation 1000*[(I_II_-(G*I_⊥_))/(I_II_+(G*I_⊥_))] with a G-factor of 0.33 for JYM20. The data was analyzed using GraphPad Prism v9.

### Isothermal titration calorimetry

ITC was performed using a MicroCal VP-ITC system (Malvern Panalytical, Westborough, MA). All proteins were dialyzed into buffer containing 150 mM NaCl, 25 mM Tris pH 8.0. Syringe containing 200 μM A-F7 was titrated into a calorimetry cell containing 20 μM SMART-MR1/5-OP-RU at 25°C. Following an initial 5 μL injection, injection volumes were 10 μL for a duration of 20 s and spaced 240 s apart to allow for a complete return to baseline. Data were processed and analyzed with Origin software. Isotherms were fit using a one-site ITC binding model with the first data point excluded from analysis. ITC was performed with one technical replicate.

### Cryo-EM sample preparation

The A-F7 TCR/SMART MR1-5-OP-RU complex was purified using SEC and best peak fraction (0.3 mg/mL) was used for grid preparation. Quantifoil R 1/2 300 Mesh, Cu grids were glow discharged at 15 mA for 60s. Prior to the grid preparation, 0.01% fluorinated octyl maltoside was added to the sample to mitigate any preferred-orientation problems. A sample volume of 3 μL was applied to the grids and subjected to vitrification using a Vitrobot Mark IV system (ThermoFisher) at 4°C with 100 % humidity. The sample was blotted for 3.0 s with a blot force of 2 using Standard Vitrobot 595 filter paper (Ted Pella, Inc) and the grids were plunge-frozen into liquid-nitrogen-cooled liquid ethane.

### Cryo-EM data collection, processing, and model building

Cryo grids of the complexes were imaged at 165,000× nominal magnification using a Falcon 4 detector (ThermoFisher) on a Glacios 2 (ThermoFisher) microscope operating at 200 kV with a calculated pixel size of 0.6975 Å. Automated image collection was performed using EPU with a nominal defocus range of –0.8 to –2.0 μm.

For A-F7 TCR/SMART MR1-5-OP-RU complex, 5,448 micrographs were collected using EPU. Data processing was carried out in CryoSPARC v4.7.1. Micrographs were aligned using Patch-motion correction and the Contrast Transfer Function (CTF)-corrected using Patch CTF Estimation in CryoSPARC. Micrographs with CTF fit worse than 10 Å were discarded. Blob picker was used to pick 2,299,717 particles from entire dataset with a minimum particle diameter of 90 Å to a maximum of 180 Å. Particles picked by blob picker were inspected and extracted with a box size of 384 pixels to yield 1,673,666 particles. A round of 2D classification was performed to select best classes (408,724 particles) that look like the desired complex. Ab-initio reconstruction was performed using these particles to get two volumes. One volume containing 248,649 particles was refined to 3.16 Å resolution map. However, this map looked distorted upon rotating by 90° and the density was fragmented in multiple regions. The other volume containing 287,776 particles yielded a nearly complete and less distorted map with 3.08 Å overall resolution, thus used for model building.

For the densities corresponding TCR and MR1 heavy chain, previously solved structure of human MAIT A-F7 TCR in complex with human MR1-5-OP-RU (PDB ID: 6PUC) without α3 domain and β_2_m was used for docking. For the density corresponding to the stabilization domain, bundle consisting of three N-terminal helices of miniaturized HLA-A*02 (PDB ID: 9NDS) was used for docking. Both models were docked in the cryo-EM density map using UCSF Chimera. The model was manually adjusted in Coot and refined using real-space refinement in PHENIX package.

## Data Availability Statement

Atomic coordinates for the SMART-MR1/A-F7 TCR complex have been deposited in the PDB under the accession code 10VM.

## Acknowledgments

This work was delivered as part of the MATCHMAKERS Team supported by the Cancer Grand Challenges partnership funded by Cancer Research UK (CGCATF-2023/100004) and the National Cancer Institute (OT2CA297575) and The Mark Foundation for Cancer Research. We acknowledge support by grant NIGMS (7R35GM125034) to N.G.S.

## Supporting Information

**Figure S1.**
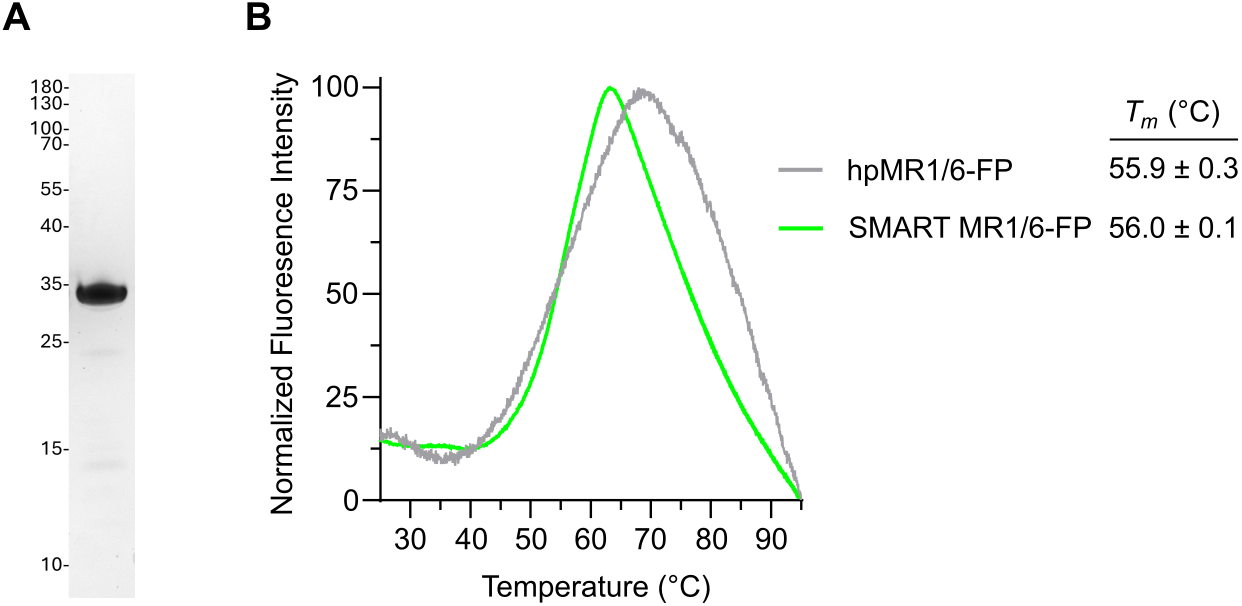
Thermal stability of SMART-MR1 compared to full-length MR1. **A**. SDS-PAGE gel of purified SMART-MR1 refolded with 6-FP. **B**. Normalized DSF traces of purified SMART-MR1 and human platform MR1 (hpMR1) refolded with 6-FP. Data are mean ± s.d. for *n* = 3 technical replicates.

**Figure S2.**
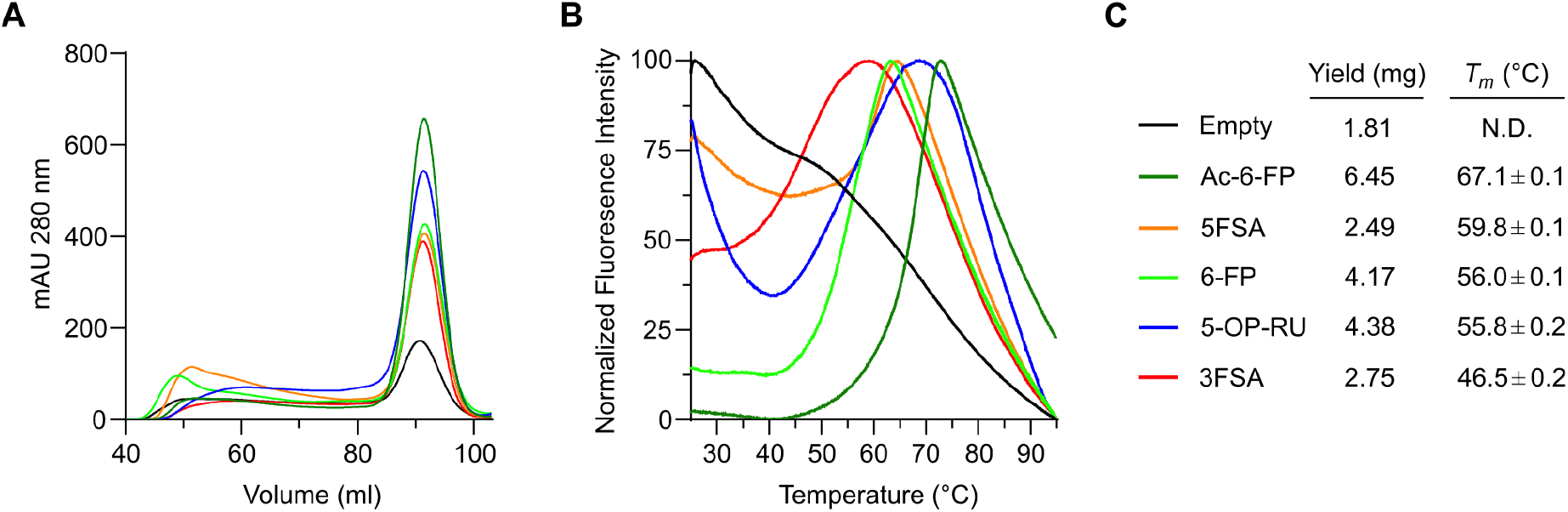
SEC traces and DSF curves of refolded SMART-MR1 complexes. **A**. Size exclusion chromatography (SEC) traces of SMART-MR1 refolded in the absence (empty) and presence of ligand. SEC traces for MR1 are color coded as shown in panel C. **B**. Normalized DSF traces of purified empty and ligand loaded SMART MR1. **C**. Summary of refolding yields obtained from SEC experiments and melting temperatures (*T*_*m*_) obtained from DSF experiments. Data are mean ± s.d. for *n* = 3 technical replicates.

**Figure S3.**
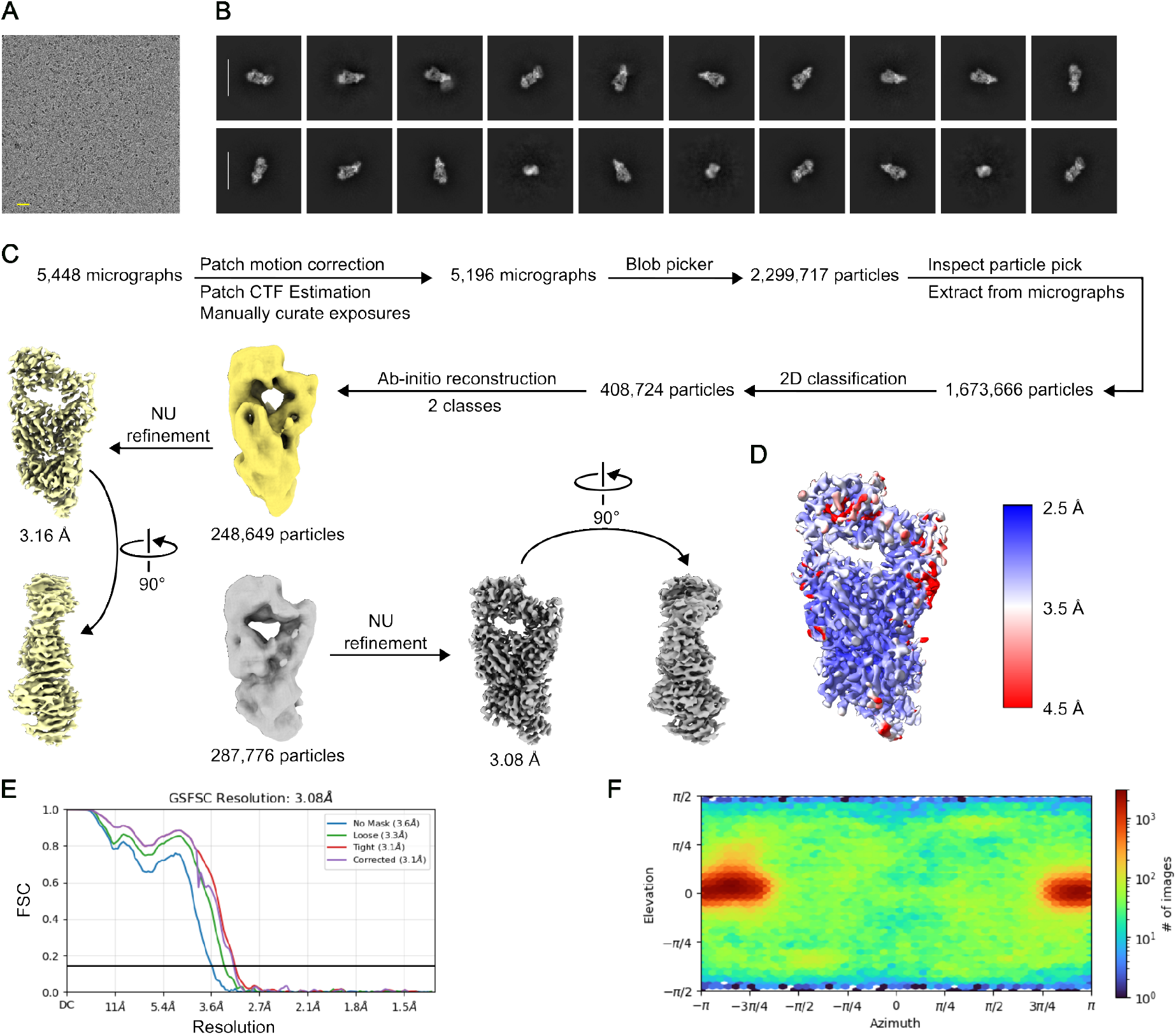
Cryo-EM data processing for A-F7 TCR in complex with SMART-MR1-5-OP-RU. **A**. Representative micrograph of the collection for A-F7 TCR/SMART-MR1-5-OP-RU complex. Scale bar in yellow is 200 Å. **B**. Representative 2D classes of A-F7 TCR/SMART-MR1-5-OP-RU complex. Scale bar in white is 150 Å. **C**. Cryo-EM data processing workflow of A-F7 TCR/SMART-MR1-5-OP-RU complex. **D**. Local-resolution estimation of reconstructed map as determined within CryoSPARC. **E**. Gold-standard FSC curves used for global-resolution estimates within CryoSPARC. **F**. Viewing direction distribution of the reconstructed map.

**Figure S4.**
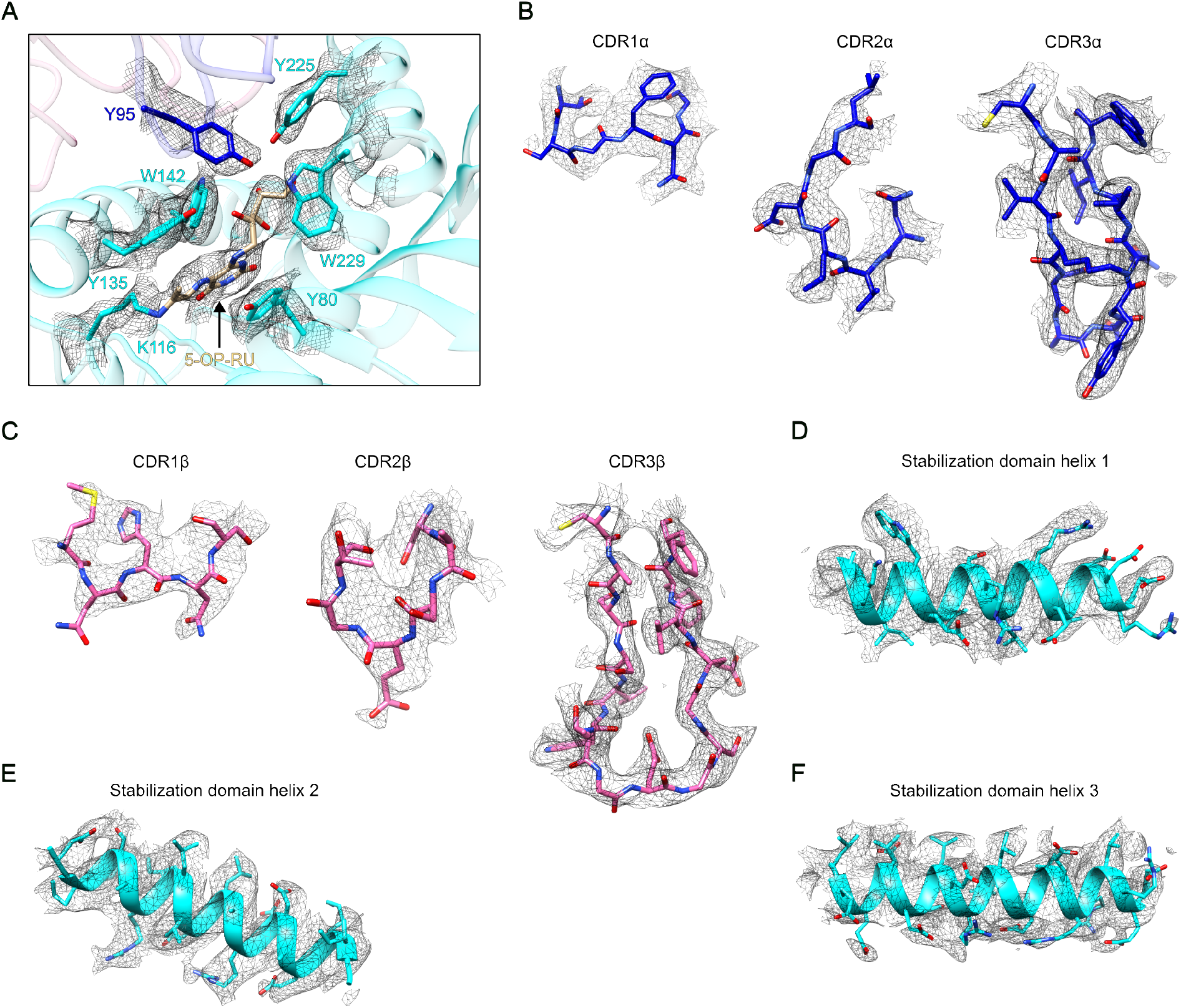
Quality of SMART-MR1-A-F7 cryo-EM map. **A**. Cryo-EM density of 5-OP-RU and the residues interacting with it. MHC is colored cyan, TCRα in blue, TCRβ in pink, and 5-OP-RU in light brown. Map is contoured at 0.196. **B**. Cryo-EM densities of CDRs of TCRα. Maps are contoured at 0.150 (CDR1α), 0.200 (CDR2α), and 0.180 (CDR3α). **C**. Cryo-EM densities of CDRs of TCRβ. Maps are contoured at 0.240 (CDR1β), 0.140 (CDR2β), and 0.120 (CDR3β). **D-F**. Cryo-EM densities of stabilization domain helices. Maps are contoured at 0.095 (D), 0.085 (E), and 0.070 (F).

**Supplementary Table S1.**
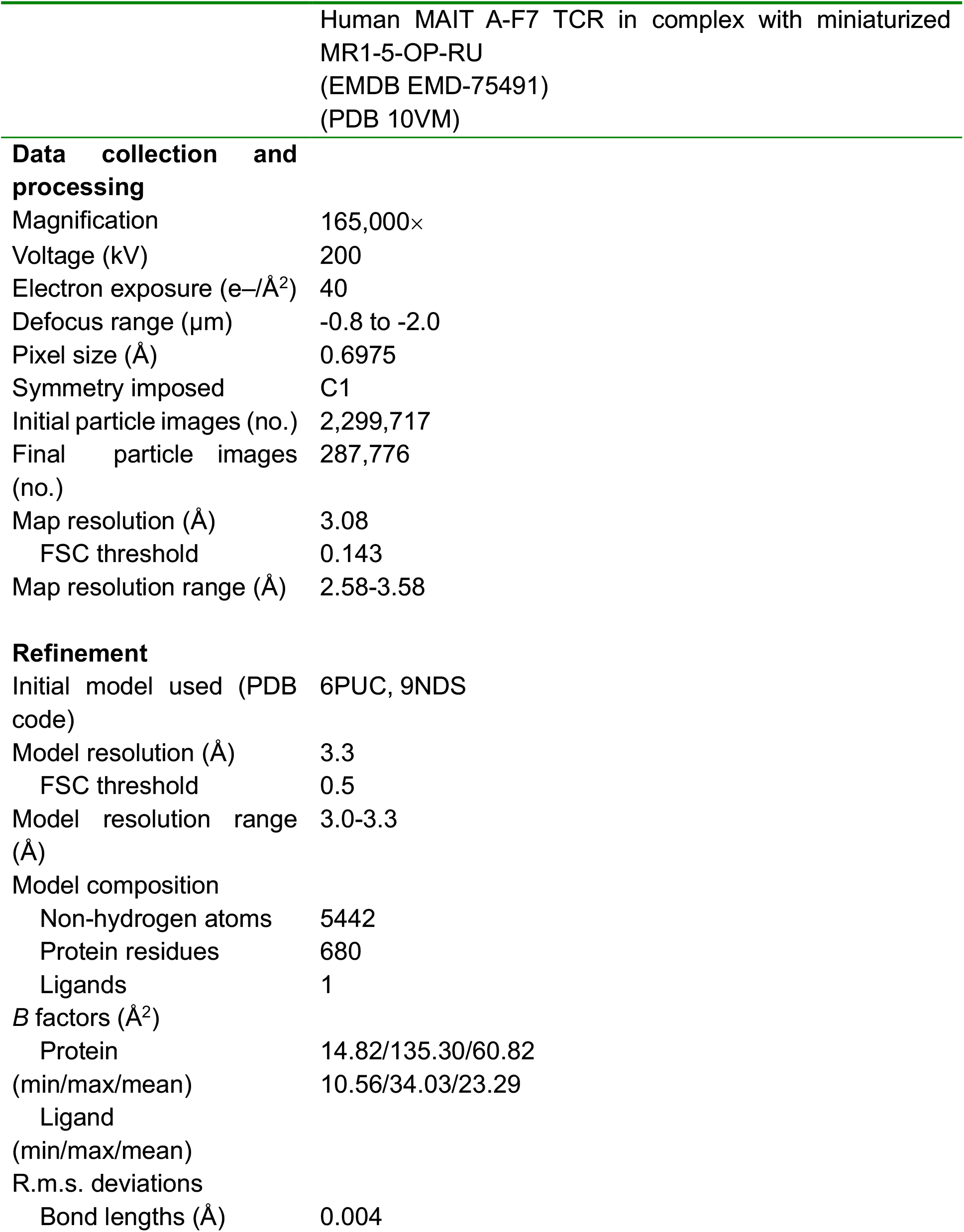
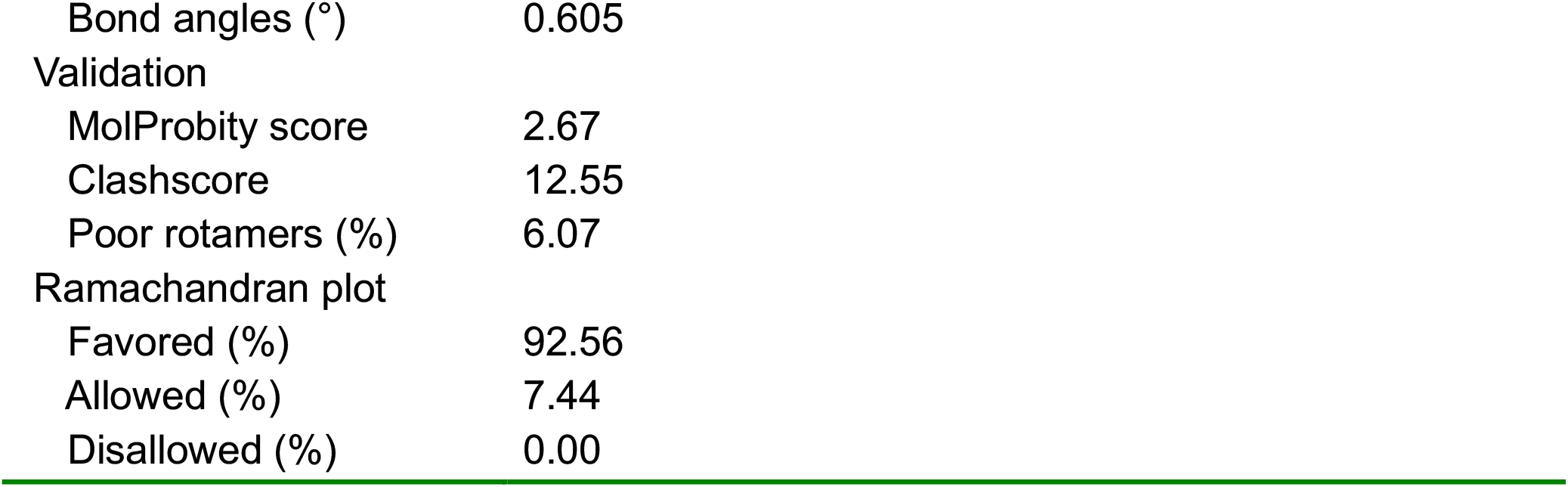
Cryo-EM data collection, refinement and validation statistics SMART-MR1/A-F7 TCR.

